# The post-transcriptional RNA splicing landscape driving oocyte division

**DOI:** 10.64898/2026.08.19.745726

**Authors:** Hady Al Shami, Andrés Gordo-Ortiz, Julia Orio-Tejada, Christelle Da Silva, Corine Blugeon, Marie-Emilie Terret, Elsa Labrune, Manuel Irimia, Adel Al Jord, Marie-Hélène Verlhac

**Affiliations:** Center for Interdisciplinary Research in Biology (CIRB), Collège de France, Université PSL, CNRS, INSERM, 75005 Paris, France; Centre for Genomic Regulation (CRG), The Barcelona Institute of Science and Technology, Dr. Aiguader 88, Barcelona 08003, Spain; Universitat Pompeu Fabra (UPF), Barcelona, Spain; GenomiqueENS, Institut de Biologie de l’ENS (IBENS), Département de biologie, École normale supérieure, CNRS, INSERM, Université PSL, 75005 Paris, France; Hospices Civils de Lyon, Inserm U1208, SBRI, 59 bd Pinel, 69500, Bron, France; Université Claude Bernard Lyon 1, Faculté de médecine Laennec, 7 rue Guillaume Paradin, 69008, Lyon, France

**Keywords:** oocyte, RNA splicing, post-transcriptional splicing, cell growth, cell division, cytoskeleton, meiosis I

## Abstract

During growth in ovaries, mammalian oocytes accumulate maternal transcripts, proteins, and metabolites that support meiotic divisions and embryogenesis. As oocytes become fully grown, RNA Polymerase II is degraded and transcription is effectively silenced. However, RNA splicing appears to be required for oocyte division, a requirement which remains poorly characterized. Here, we define this post-transcriptional RNA splicing landscape of fully grown mouse oocytes and develop a computational resource that identifies post-transcriptional splicing events, signatures, sequence features, and protein binding motifs responsive to splicing perturbation. We show that, unlike transcription, RNA splicing in fully grown oocytes is essential for meiotic progression. Applying our resource across perturbations, we identify hundreds of genes whose proper splicing is required for oocyte division and reveal that distinct perturbations converge on overlapping regulatory programs controlling cytoskeletal organization and cell cycle progression that drive oocyte division. We experimentally validate selected resource-derived predictions using chemical and protein-based splicing perturbations. These findings demonstrate that extensive RNA processing persists after transcription has ceased and position post-transcriptional splicing as a fundamental regulator of mammalian oocyte development.

## Introduction

Female fertility is at risk in modern societies, with poor oocyte quality being a major causal factor. Oocytes are exceptionally large cells whose quality and developmental potential depend heavily on the successful accumulation of maternal proteins and messenger RNAs (mRNA) during their growth in the ovary^1^. When Prophase I oocytes reach their fully grown stage in mice and humans, transcriptional activity drops drastically up to 10 fold^2–13^. This decrease in transcription is caused by the controlled degradation of RNA Polymerase II in mouse oocytes, with evidence suggesting comparable RNA Polymerase II degradation in fully grown human oocytes^5^. Despite RNA Polymerase II degradation causing a transcriptional halt, the splicing of RNA is nonetheless required for fully grown mouse oocytes to successfully divide when they resume meiosis^4^, strongly arguing for critical post-transcriptional RNA splicing activity in these cells.

RNA splicing has often been viewed to mostly occur co-transcriptionally^14^, yet post-transcriptional splicing is increasingly appreciated as a key regulator of gene expression across health and disease contexts^15,16^. The timing of splicing substantially influences alternative splicing efficiency, impacting the combinations of exons and/or introns eventually kept in the final mature mRNA. The relative contributions of co-transcriptional and post-transcriptional splicing to splicing regulation, and their respective biological impacts, constitute an active area of research^15^.

Post-transcriptional splicing regulation as well as cell division control across cell types depend on nuclear speckles, essential nuclear biomolecular condensates^4,17–27^. In mouse oocytes, nuclear speckles evolve in size and number during growth^4,28^, and our new observations suggest that they also reorganize during human oocyte growth (Fig. S1A-E). As mouse oocytes become fully grown, splicing activity becomes concentrated primarily in nuclear speckles^4^, and the perturbation of speckle organization is linked to defects in RNA splicing and oocyte division^4^. Together, these observations prompted us to precisely investigate the importance of post-transcriptional splicing in fully grown oocytes, and its influence on the subsequent oocyte division that should produce a fertilizable egg.

Here, we defined the post-transcriptional RNA splicing landscape of fully grown mouse oocytes and examined its impact on meiotic cell division using chemical perturbations, deep RNA sequencing, computational analyses, and live microscopy. We created a computational resource (*<u>Oocyte Splicing</u> <u>Explorer</u>*), which enabled us to explore and determine post-transcriptional pre-mRNA splicing events, splicing signatures, sequence features, and protein binding motifs that respond to splicing perturbation in grown oocytes. We found that transcription in fully grown oocytes is dispensable for meiotic division, whereas spliceosome activity remains essential for the meiotic development of oocytes. By applying this resource across multiple splicing perturbations, we identified hundreds of genes whose correct splicing is essential for oocyte division and showed that distinct perturbations converge on shared regulatory programs underpinning oocyte division like that of the cytoskeleton, the cell cycle and cytokinesis. Finally, we experimentally validated selected predictions generated by the resource using chemical splicing modulators and candidate splicing factor overexpression-based perturbations. Altogether, our work shows that developmental progression can rely on extensive RNA processing independently of ongoing transcription and introduces a broadly useful resource of key candidate genes whose splicing is essential for oocyte division that fosters female fertility. Beyond reproduction, our resource is relevant for cancer biology since oocytes share cell division-related properties with cancer cells^29^, which in turn often harbor perturbed splicing patterns responsible for enhanced tumorigenesis^16,30,31^.

## Results

### Transcription in fully grown oocytes is dispensable for meiotic progression and division

Discoveries from decades ago implied that oocytes from many species, including humans, shut transcription down at the end of their growth^6–13,32^. However, the impact of residual transcription on the developmental potential of fully grown mammalian oocytes remained poorly characterized. Conflicting results were reported: bovine oocytes allowed to resume meiosis in the presence of the reversible transcription inhibitor DRB exhibited impaired progression^33^, whereas another study using the same drug at similar concentrations in mouse oocytes reported the opposite effect^34^. No study, however, has examined the consequences of inhibiting transcription in fully grown oocytes and subsequently assessing the impact on meiotic progression. To thus investigate the importance of residual transcription on fully grown mouse oocyte division, we acutely treated them in Prophase I with the reversible transcription inhibitor DRB^34,35^ for 5 hours before washing out the inhibitor and allowing oocytes to resume meiosis (Fig. 1A). DRB treatment prior to meiosis resumption had no discernable impact on the first meiotic division of oocytes. The treated oocytes resumed meiosis, indicated by Nuclear Envelope Break Down (NEBD), and divided, evidenced by first polar body extrusion (PBE), with identical timings and efficiencies as control oocytes (Fig. 1B). Similarly, α-amanitin, an irreversible transcription inhibitor, had no impact on meiotic progression in oocytes (Fig. S2).

**Figure 1.**
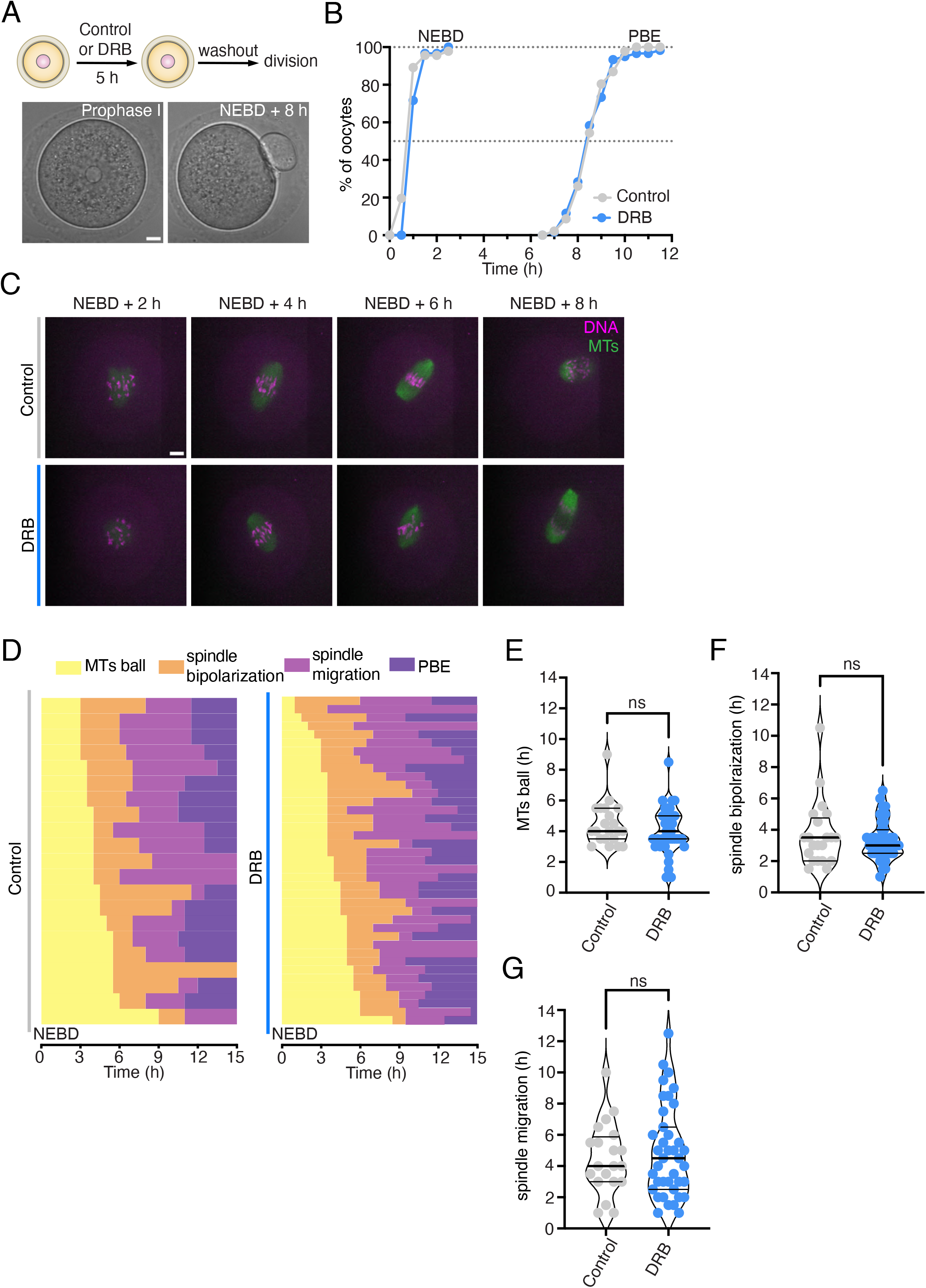
Transcription inhibition in oocytes does not affect meiotic progression and division. **A.** Schematic representation of the experimental approach (up). The lower images correspond to transmitted light images of a fully grown oocyte in Prophase I (left) and after its first polar body extrusion (right; 8h after Nuclear Enveloppe BreakDown, NEBD). Oocyte are maintained arrested in Prophase I for 5h thanks to the presence of Milrinone in the culture medium. The washout corresponds to removal of all drugs, Milrinone included. **B.** Percentag e of control and DRB-treated oocytes that undergo NEBD and extrude their first polar body (PBE) as a function of time (h) after washout from drugs. The 0h time point marks the first time point after washout. N= 46 oocytes for control; N= 59 oocytes for DRB; 3 independent experiments. **C.** Control (top) and DRB-treated (bottom) oocytes montage of a time-lapse movie where chromosomes (pink) and microtubules (green) are fluorescently labelled. **D.** Stacked bar chart showing the timing of the different phases of the first meiotic division in control and DRB-treated oocytes: microtubule ball (yellow), spindle bipolarization (orange), spindle migration (pink), and polar body extrusion (purple). Each horizontal bar corresponds to a single oocyte. The 0h time point marks the first time point after washout. N= 21 oocytes for control; N= 39 oocytes for DRB; 2 independent experiments. **E, F, G.** Respective quantification of the durations (h) of the microtubule ball phase, the spindle bipolarization phase, and the spindle migration phase in control and DRB-treated oocytes. Data are shown as mean ± SD. P values were calculated using Mann-Whitney test. Scale bars are 10 µm. ns = not significant; * = p<0.05; ** = p<0.01; *** = p<0.001; **** = p<0.0001.

To further evaluate the effect the DRB treatment, we examined distinct sequential steps of cell division during meiosis I in oocytes. First, oocytes, due to lack of canonical centrosomes, assemble their meiotic spindle around condensed chromosomes in the form of a microtubule ball^36–38^. This microtubule ball then progressively transforms into a bipolar structure via an auto-assembly process a few hours after entry into meiosis^39^. Once the spindle axis is formed, the spindle migrates along its long axis to the closest cortex due in part to a dynamic F-actin meshwork^40–43^. Finally, anaphase I takes place around 8h after NEBD followed by the extrusion of the first polar body. In control and DRB-treated oocytes with labeled chromosomes and microtubules (Methods), we precisely captured all these events in live (Fig. 1C; Movies S1 and S2) and measured the duration of each specific stage as shown in Fig. 1D. We did not observe any significant difference between controls and DRB treated oocytes in terms of the duration of each of these key stages (Fig. 1E, F, G). As the proper duration of key spindle assembly stages is essential to oocyte quality^39,44^, our data are consistent with recent findings^5^ and indicate that residual transcription in fully grown oocytes plays no major role in their developmental potential.

### Acute treatment with splicing modulators induces splicing alterations in fully grown oocytes

Unlike transcription, RNA splicing in fully grown oocytes plays a key yet poorly characterized role in their developmental potential^4^. To gain a wide range of molecular insights into the importance of RNA splicing in fully grown oocytes, we developed a computational resource based on RNA sequencing of fully grown oocytes treated independently with three established splicing modulators (Fig.2A; Resource: *<u>Oocyte Splicing Explorer</u>*). We incubated oocytes with either Tubercidin, an adenosine analogue shown to disrupt nuclear speckles and alter splicing in cancer cells and oocytes^4,30,45,46^, Pladienolide B or Spliceostatin A, two distinct splicing modulators that specifically target the SF3B complex of the spliceosome^47–51^. After 5-hour incubations with the compounds, we extracted the RNA and prepared libraries for deep poly-A bulk RNA sequencing – yielding an average of 360 million reads per condition (Fig. 2A and Methods). To assess the impact of these three chemicals on the oocyte transcriptome, we performed bioinformatical analyses of gene expression and RNA splicing for each data set (Fig. 2A-C; Supplementary Tables 1 and 2). We found that these acute incubations with Pladienolide B and Spliceostatin A treatments led to minimal changes in gene expression (7 and 3 genes respectively), whereas Tubercidin did not cause any detectable gene expression alterations (Fig. 2B; (|log2FC| ≥ 1.5, FDR ≤ 0.05)). The incubations, however, were sufficient to cause measurable and significant splicing alterations (Fig. 2C). To proceed, we leveraged VAST-TOOLS to align our samples and quantify splicing events from the curated and annotated database VastDB^52^, and combined it with the betAS^53^ package which calculates statistical significance across conditions by modelling thousands of simulations from the beta distribution. As previously done^54^, we defined a splicing event as significant when it reached an absolute difference in its spliced-in rate, known as percent spliced-in or PSI, of at least 10% (|ΔPSI| ≥0.1, False Positive Rate (FPR) ≤0.05). We detected thousands of significant splicing events across treatments (Fig. 2C; Fig. S3A), including exons, introns, alternative donor sites (Alt5) and alternative acceptor sites (Alt3). All three treatments induced comparable numbers of splicing pattern alterations, both in total and when broken down by event type (Fig. 2D). When splicing events were grouped by their corresponding genes, the three treatments also showed comparable total numbers of spliced genes, yet with some limited overlap between uniquely spliced genes (Fig. 2E). Thus, short incubations of fully grown oocytes with mechanistically distinct RNA splicing modulators does not significantly change gene expression but does induce extensive alterations in RNA splicing.

**Figure 2.**
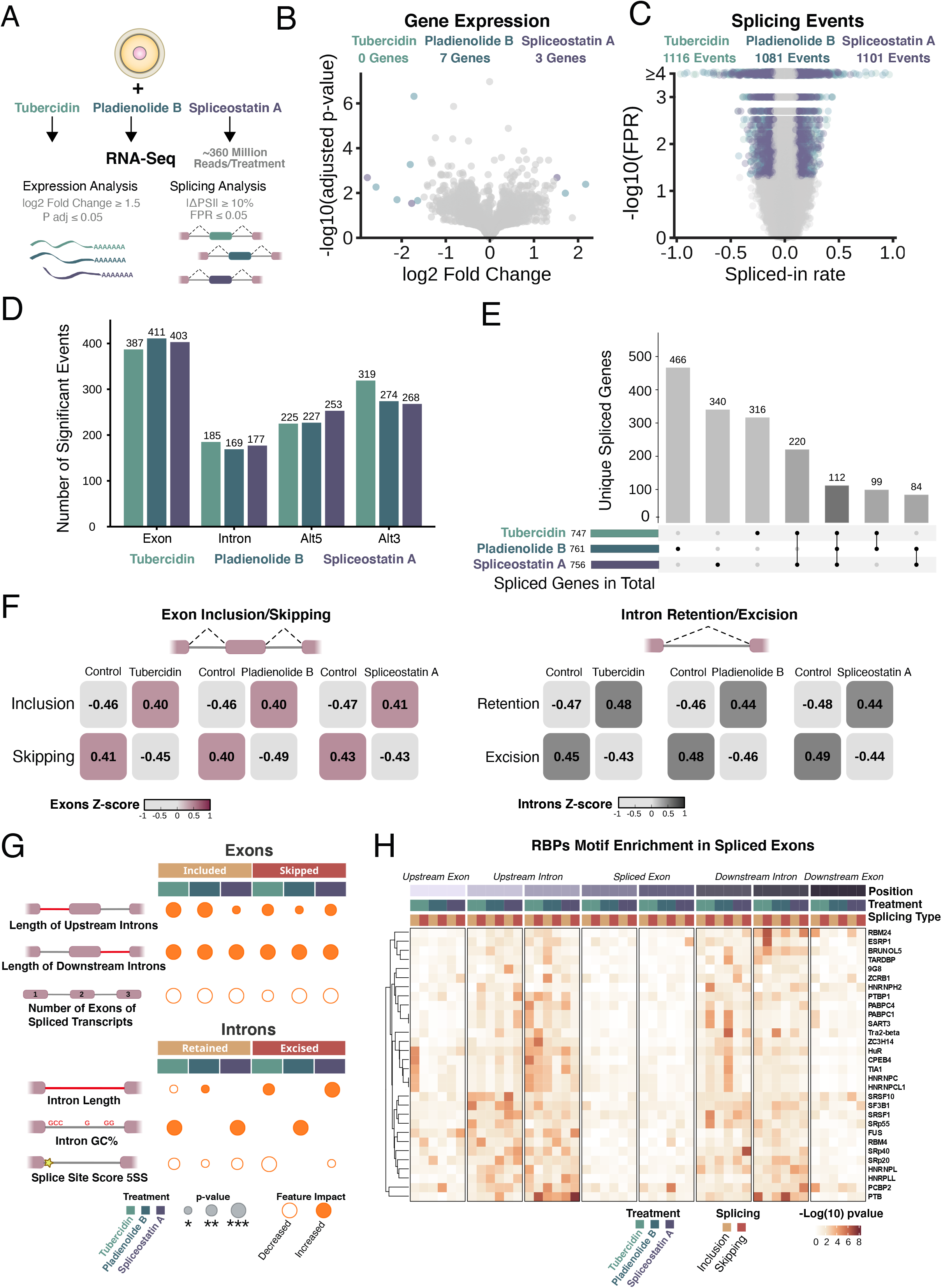
Acute splicing perturbations in oocytes impact RNA splicing but not gene expression. **A.** Schematic of the experimental design. Mouse oocytes, maintained arrested in Prophase I, were treated with either Tubercidin, Pladienolide B or Spliceostatin A for 5h. RNA was extracted from 3 independent biological replicates of each treatment for sequencing, yielding an average of 360 million reads per condition. **B.** Volcano plot showing the sequenced genes by their log2 fold change (Treatment/Control) and –log10 adjusted p-value. Differentially expressed genes reach a log2 fold change of at least 1.5 and an adjusted p-value lower than 0.05, in which case they are colored by the corresponding treatment. **C.** Volcano plot showing splicing events by their spliced-in rate, also known as PSI, and the False Positive Rate (FPR, see Methods). Significant splicing events reach a |ΔPSI| of at least 0.1 (10%) and a FPR lower than 0.05, in which case they are colored by the corresponding treatment. **D.** Bar plots depicting the total number of significant splicing events per type (either cassette exons, introns, alternative donors/5’ or alternative acceptors/3’) and treatment. **E.** The horizontal bars depict the total number of genes being significantly spliced across treatments, whereas the vertical bars show the intersection of those genes, either unique or shared, between treatments. **F.** Chequerboard plot showing the differences in the amount of included and skipped exons (left), or retained and excised introns (right). Each small tile compares the mean PSI for Control versus Treatment samples for a collapsed cluster of differential events; values printed in the tiles are the collapsed mean scaled PSI (median-centred, MAD scaled and truncated to the −1 to +1 range). Patterns were defined by hierarchical clustering (Spearman correlation). Only significant events were included (FPR≤0.05 and |ΔPSI|≥0.1). **G.** Summarizing table of the significant features found in spliced exons, introns and neighboring sequences, per treatment, and compared to a background group of non-spliced events. Exons and introns are respectively categorized as included and retained (ΔPSI > 0.1) or skipped and excised (ΔPSI < -0.1). Statistical pairwise comparisons were performed using Mann-Whitney tests (*P ≤ 0.05, **P ≤ 0.01, ***P ≤ 0.001, ****P ≤ 0.0001). **H.** Comprehensive RBP motif enrichment analysis across treatments of the significant exons or neighboring sequences. The heatmap shows the highest -log10 (p-values) for the top RBPs, sorted by maximum significance across treatments. Rows are clustered by Euclidean distance; columns are ordered by genomic position, drug and splicing direction (Inclusion, Skipping). The positions correspond, from left to right (5’—>3’), to the last 50bp of the upstream exon, the first and last 250bp of the upstream intron, the first and last 50bp of the spliced exon and likewise for the downstream intron and downstream exon.

### Overlapping RNA splicing events, signatures, sequence features and motifs revealed by splicing modulators

To gain further insight into the impact of the treatments on splicing patterns, we examined how the three treatments affected the balance of exon inclusion-skipping and intron retention-excision. All three compounds surprisingly produced a similar pattern: comparable amounts of exons showed an enhanced inclusion *versus* skipping, and comparable numbers of introns exhibited an increased retention *versus* excision (Fig. 2F; Fig. S3B-C). Both event types showed signs of a symmetrical distribution of splicing after 5 hours of treatment with the modulators. Thus, the induction of extensive RNA splicing alterations via diverse chemical modulators in oocytes produces seemingly similar qualitative outcomes in terms of splicing patterns. Therefore, both event types showed signs of a symmetrical distribution of splicing after 5 hours of treatment with the modulators, while general splicing inhibitors usually lead to increased exon skipping and intron retention^55^.

Since transcript sequence landscapes and RNA-protein binding site motifs are well described modifiers of splicing patterns, we documented multiple sequence-specific structural features and RNA-binding protein (RBP) motif enrichments in our lists of transcripts (Fig. 2G-H; Fig. S4, S5, and S6). First, we characterized sequence-specific features enriched in the significantly spliced exons, introns, or neighboring sequences (Fig. 2G; Fig. S4 and S5). To proceed, we used Matt^56^– a toolkit for exon and intron feature extraction and comparison. Matt computes splice-site strengths using published maximum entropy models, predicts branch points, or quantifies the length of the pyrimidine tract, among other analyses (Supplementary Data 1). Significantly spliced exons exhibited common features across treatments, such as being surrounded by longer adjacent introns, or belonging to transcripts with fewer exons in average (Fig. 2G, Fig. S4A-C). These features are consistent with a broader shift from intron definition to exon definition, in which splice-site recognition across introns breaks down once introns become sufficiently long, allowing exon-based recognition to take over, as suggested by several studies^57–59^. Furthermore, a recent hallmark study demonstrated that exons flanked by longer introns – a signature of exon definition – are indeed more sensitive to spliceosome perturbation, although the authors examined genetic rather than chemical perturbations^60^. The different treatments presented some disparities as well (Fig. S4D-F). For instance, with Tubercidin and Spliceostatin A, included exons harbored weaker 3’ splice sites, whereas skipped exons harbored weaker 3’ splice sites and longer polypyrimidine tracts in their downstream introns (Fig. S4E-F). On the other hand, Pladienolide B-included exons had fewer predicted branch points in their upstream intron, a feature not detected with the two treatments (Fig. S4D). In parallel, significantly spliced introns exhibited a more nuanced behavior, depending on retention *vs.* excision status or the splicing modulator used (Fig. 2G; Fig. S5A-F). For example, Tubercidin- and Spliceostatin A-excised introns were longer but Pladienolide B-retained introns were the longer ones. Also, Pladienolide B-excised introns had higher GC content, whereas Tubercidin- and Spliceostatin A-retained introns were the ones to show higher GC content (Fig. 2G, Fig. S5A-B). Other features showcased a more uniform interpretation in all spliced introns across treatments, such as a weaker 5’ splice site, differential transcript length, or higher GC contents of the exons neighboring the spliced introns (Fig. S5C-F). Together, these data suggest that the splicing alterations preferentially affect exons and introns with specific structural features – such as longer neighboring introns, weaker splice-site signals, or distinct GC content patterns – that are known to influence splice-site recognition^55,61–63^. These characteristics may make certain regions more responsive to spliceosome perturbation, delivering a plausible explanation for the consistent types of splicing changes observed across treatments.

Second, we investigated whether the spliced sequences significantly impacted by the treatments were enriched in motifs known to be bound by certain RBPs (Fig. 2H, Fig. S6A-E). For this analysis, we chose rMAPS2^64^ – a computational motif enrichment analysis tool for experimentally assessed RBPs. The results on the significantly spliced exons or introns revealed a myriad of motif enrichments comprising RBPs from the core spliceosome-associated machinery like U2AF2, involved in 3’ splice site recognition, and SF3B1, a direct target of Spliceostatin A and Pladienolide B^65^, as well as key splicing regulators like SRSF proteins^30^ (Fig. 2H, Figs. S6A and C-D; Supplementary Data 2). In included and skipped exons, motifs were enriched predominantly in the adjacent introns (Fig. 2H, Fig. S6A-B), whereas retained introns showed increased enrichment within the intronic sequence (Fig. S6C-D). Another difference lied in the total number of enriched motifs: while included and skipped exons exhibited comparable numbers (Fig. S6B), spliced introns had significant motif enrichment differences when comparing retained and excised (Fig. S6E). These findings are consistent with the known mechanisms of splicing modulators, which interfere with the recruitment of splicing regulators in a context-dependent manner. While SF3B inhibitors target the U2 snRNP by binding the branch point adenosine pocket, their differential splice site sensitivities are primarily dictated by sequence-dependent features. Furthermore, the downstream interference of splicing modulators with other splicing factors at multiple stages of splicing, both *in vitro* and *in vivo*^66^, suggests that these modulators can alter the regulatory interplay of a myriad of cell-type and context-dependent splicing factors. Overall, our collective analyses indicate that treating fully grown oocytes with splicing modulators leads to significant and specific splicing alterations: they affect multiple types of splicing events, involve sequence features that influence splice-site recognition, and are enriched for motifs of splicing- and spliceosome-linked RBPs.

### Splicing modulators impact genes involved in meiotic progression and division of oocytes

To evaluate computationally the functional impact of the induced splicing alterations, we made use of VastDB’s predicted annotations of splicing events which inform on whether an event will disrupt the coding sequence by shifting the open reading frame (ORF), concur in an alternative isoform, or impact regulatory regions (Fig. 3A; Supplementary Table 3). When uploading significant splicing events induced by the splicing modulators, all three treatments showed similar proportions of impact annotations - with the majority belonging to alternative isoforms (46.1% to 49.2%), followed by events disrupting the ORF (23% to 26.6%) and changes in the 5’ untranslated region (UTR; 17.9% to 18.3%). This suggests that the detected splicing alterations may diversify isoform production, affect their regulation, and fine-tune protein production. To finally capture the gene ontology of these differentially spliced events, we examined enrichment maps from across Gene Ontology categories (*i.e. biological process, cellular component, molecular function*; Supplementary Table 4). The analyses revealed that a majority of the common terms across the three treatments were related to processes implicated in cell division (Fig. 3B). Thus, and although certain treatments can affect a partially distinct set of genes (Fig. 2E), all converge on gene ontology categories linked to cell division processes (Fig. 3B), implying that splicing in fully grown oocytes consistently impacts pathways central to meiotic progression and division.

**Figure 3.**
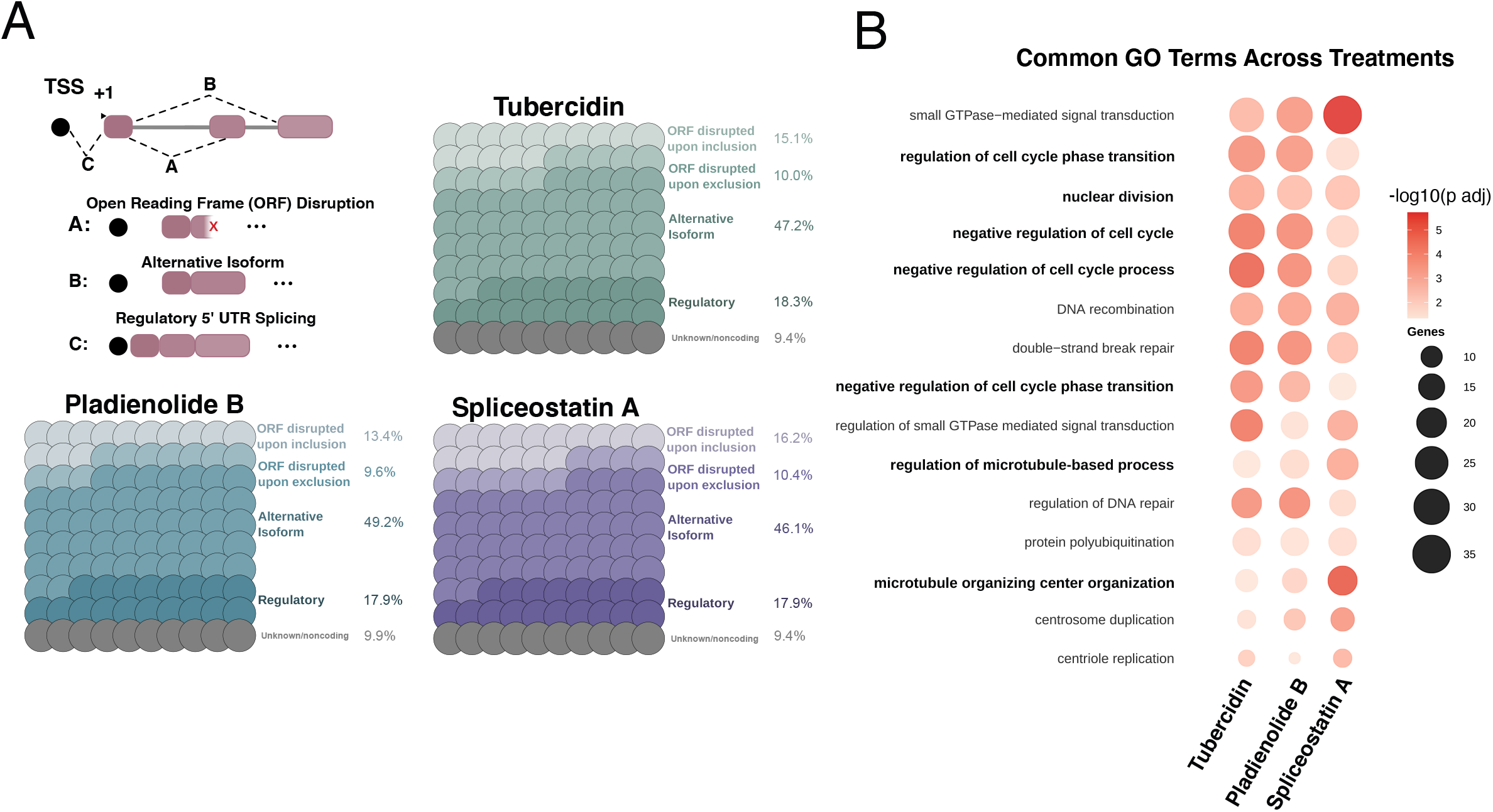
Compound-driven changes in splicing are predicted to disrupt meiotic progression and cell division-related processes. **A.** Proportions of the different functional impact annotations derived from the significant splicing events across treatments according to VastDB. The schematic displays the three types of functional consequences analyzed: open-reading frame (ORF) disruption, an alternative complete isoform, or regulatory impact in the untranslated region (UTR). **B.** Enrichment map for shared GO Biological Process terms across treatments of significantly spliced genes. The node size is proportional to the number of genes associated with the GO term and the colour intensity to the adjusted p-value of the enrichment. Note that cell division-related processes are in bold.

### Experiments validate resource-derived predictions by showing that splicing perturbations in fully grown oocytes hinder meiotic progression and division

We next sought to experimentally validate selected computational predictions using chemical modulators and resource-derived candidate protein perturbations (Fig. 4). Our analyses predicted splicing alterations in genes functionally associated with meiotic progression and division (Fig. 3B; Supplementary Table 4). For instance, top hits included regulators of cell cycle progression, microtubules and their organizing centers, essential for meiotic spindle dynamics, as well as small GTPases, involved in the regulation of F-actin, DNA, spindle, and polar body dynamics^39,67^. To test these predictions, we first probed experimentally the impact of two splicing modulators - Tubercidin and Spliceostatin A - on meiotic progression and division of oocytes. We washed out the pharmacological agents after a 5-hour incubation with fully grown oocytes and monitored their progression through meiotic division.

**Figure 4.**
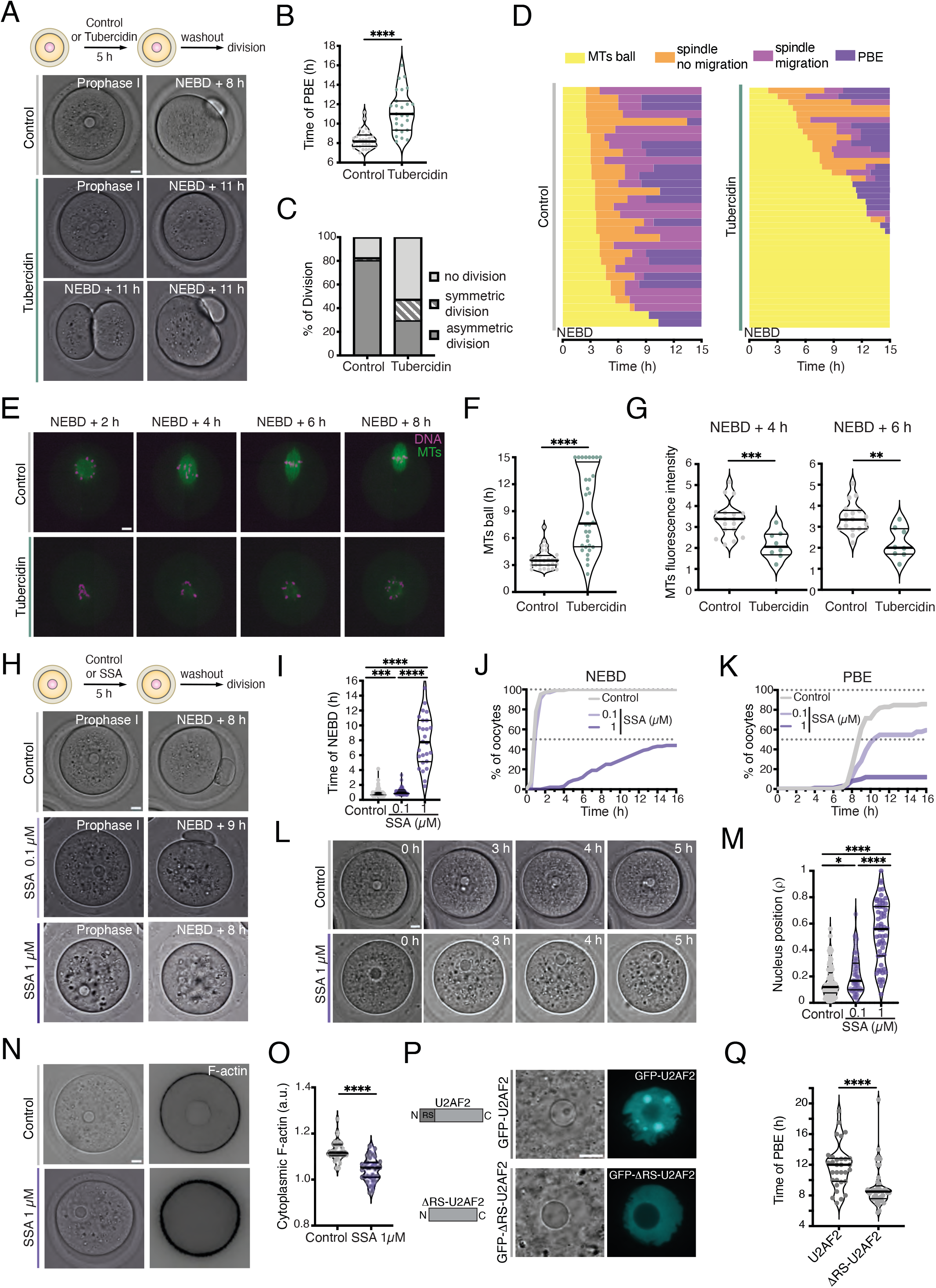
Experimental validations reveal that splicing perturbations in fully grown oocytes hinder meiotic progression and division. **A.** Schematic representation of the experimental approach (top panel). The central images correspond to transmitted light images of a fully grown control oocyte in Prophase I and after its first polar body extrusion (8h after Nuclear Envelope Break Down, NEBD). Oocyte were maintained arrested in Prophase I for 5h thanks to the presence of Milrinone in the culture medium. The washout corresponds to removal of all drugs, Milrinone included. The lower images show transmitted light pictures of Tubercidin-treated oocytes with Prophase I oocyte and oocytes 11h after NEBD either undivided, symmetrically or asymmetrically divided. **B, C.** Respective quantification of the timing of first PBE and of the percentages of the types of division in control and Tubercidin-treated oocytes. N= 57 oocytes for control, N= 57 oocytes for Tubercidin, 3 independent experiments. **D.** Stacked bar chart showing the timing (h) of the different phases of the first meiotic division in control and Tubercidin-treated oocytes: microtubule ball (yellow), spindle bipolarization (orange), spindle migration (pink), and polar body extrusion (purple). Each horizontal bar corresponds to a single oocyte. N= 29 oocyte for control, N= 32 oocytes for Tubercidin, 2 independent experiments. **E.** Control (top) and Tubercidin-treated (bottom) oocytes montage of a time-lapse movie where chromosomes (pink) and microtubules (green) are fluorescently labelled. Scale bar 10 µm. **F.** Quantification of the duration (h) of the microtubule ball in control and Tubercidin-treated oocytes. Data are shown as mean ± SD. P values were calculated using Mann-Whitney test. **** = p<0.0001. **G.** Respective quantification of microtubule fluorescence intensities at 4 and 6 h after NEBD in control and Tubercidin-treated oocytes. N= 19 oocytes for control, N= 8 oocytes for Tubercidin, 2 independent experiments. Data are shown as violin plots with median ± quartiles. P values were calculated using Mann-Whitney test. ** = p<0.01; *** = p<0.001 **H.** Schematic representation of the experimental approach (top panel). The upper images correspond to transmitted light images of a fully grown oocyte in Prophase I (left) and after its first polar body extrusion (right; 8h after Nuclear Envelope Break Down, NEBD). Oocyte are maintained arrested in Prophase I for 5h thanks to the presence of Milrinone in the culture medium. The washout corresponds to removal of all drugs, Milrinone included. The central and lower panels show examples of delayed PBE in Spliceostatin (SSA)-treated oocytes at different doses. Scale bar 10 μm. **I.** Quantification of the time of NEBD in control and SSA-treated oocytes at 0.1 and 1 µM. **J, K.** Percentage of control and SSA-treated oocytes that undergo NEBD and extrude their first polar body (PBE) as a function of time (h) after washout from drugs. N= 88 oocytes for control, N= 64 oocytes for SSA at 0.1 µM; N= 64 oocytes for SSA at 1 µM; 6 independent experiments. **L.** Nucleus drifts to the cortex after SSA treatment at 1 µM. Control (top) and SSA-treated (bottom) oocyte montages of transmitted light movies in Milrinone. Scale bar 10 μm. **M.** Quantification of nucleus position (r = d (nucleus center - oocyte center)/(oocyte radius - nucleus radius) in control and SSA-treated oocytes at two doses (0.1 and 1 µM). r varies between 0 (perfectly centered nucleus) and 1 (completely off-centered nucleus). **N.** SSA treatment at 1 µM impacts F-actin. Transmitted light images (left) and selected Z-plane of cytoplasmic F-actin labelling of control (top) and SSA 1 µM-treated (bottom) oocytes after 5 hours of treatment. Scale bar 10 μm. **O.** Normalized cytoplasmic F-actin intensity of control and SSA 1 µM-treated oocytes after 5 hours of treatment. N= 30 oocytes for control; N= 33 oocytes for SSA at 1 µM; 3 independent experiments. Data are shown as violin plots with median ± quartiles. P values were calculated using Mann-Whitney test. * = p<0.05; *** = p<0.001; **** = p<0.0001. **P.** Expression of wild type and mutant U2AF2 tagged with GFP in Prophase I oocytes. Transmitted light images (left) and Fluorescence images (right) of oocyte nuclei over-expressing either wild type U2AF2 (upper panel) or mutant ΔRS-U2AF2 (lower panel). Note that only wild type U2AF2 incorporates into round structures that evoke nuclear speckles. **Q.** Quantification of the time in hours of polar body extrusion of oocytes expressing either wild-type U2AF2 and ΔRS-U2AF2. N= 36 oocytes for wild-type U2AF2, N= 47 oocytes for ΔRS-U2AF2, 3 independent experiments. Data are shown as violin plots with median ± quartiles. P values were calculated using Mann-Whitney test. Scale bars are 10 µm. **** = p<0.0001.

First, in Tubercidin’s case, we found that the treatment delayed the overall cell cycle progression (Fig. 4A-B) and caused a net reduction in successful division (Fig. 4C). After this general overview of the induced phenotype, we probed further to find that Tubercidin significantly perturbed the asymmetry of the first division, inducing about 20% of symmetric cleavages (Fig. 4 A and C). By live-monitoring microtubules and chromosomes after Tubercidin treatments, we observed a significant delay in spindle bipolarization (Fig. 4D-E; Movie S3 and S4) with a persistent microtubule ball (Fig. 4F) and a failure of chromosomes to form correctly aligned metaphase plates (Fig. 4E). In the absence of canonical centrosomes defining spindle poles at mitosis entry, the capacity of the microtubule ball to form a bipolar spindle depends on microtubule nucleating and stabilizing factors^37–39,68^ as well as on the presence of microtubule motors, such as Eg5 or HSET^36,69,70^. A more precise follow-up of spindle assembly showed that the microtubule ball displayed a persistent reduction of microtubule density after treatment with Tubercidin when compared to controls (Fig. 4G; Movie S5 and S6), an observation consistent with an extended duration of the microtubule ball stage (Fig. 4F). These findings show that a significant delay in spindle assembly underlies the observed delay in polar body extrusion.

Second, in Spliceostatin A’s (SSA) case, the treatment had strong downstream consequences on cell cycle progression (Fig. 4H). We found a significant dose-dependent delay in meiosis resumption (Fig. 4H-J), which decreased the percentage of oocytes undergoing NEBD (Fig. 4J) as well as those being able to extrude a first polar body (Fig. 4K). We also made an unexpected observation of the dose-dependent drifting of the nucleus from the cell center to the cortex, visible as of 3 hours post-SSA incubation (Fig. 4L-M; Movie S7 and S8). Since nucleus positioning depends on cytoplasmic F-actin^4,71,72^, we thus assessed F-actin organization in oocytes treated with higher doses of SSA. We detected a significant reduction in the intensity of F-actin inside the cytoplasm of oocytes treated for 5 hours with SSA (Fig. 4N-O), which may be the causal driver of this newly documented phenotype. Collectively, these experimental findings imply that splicing in fully grown oocytes is essential for the correct deployment of multiple key drivers of meiotic division. The experiments also show that broad splicing modulation in fully grown oocytes causes multiple defects in the control of oocyte division, thereby validating our resource-based predictions in oocytes.

Finally, and to further validate our resource-derived analyses, we performed protein-based perturbation experiments targeting a candidate splicing factor. Our computational analyses identified a significant enrichment in binding sites for U2AF2 in the intron retention events observed upon treatment with Tubercidin and Spliceostatin A (Fig. S6D), suggesting that this protein plays a role in the post-transcriptional splicing landscape of fully grown oocytes. U2AF2 is key splicing factor that helps the binding of U2 small nuclear riboprotein to the pre-mRNA branch point during early steps of the splicing reaction^30,73–76^, and overexpression of the wild type U2AF2 interferes with and alters splicing activity in human cells^77^. We therefore overexpressed the wild type U2AF2, by injecting large doses of coding RNA into fully grown oocytes, as well as a mutant form lacking its N-terminal RS domain (ΔRS-U2AF2) and shown *in vitro* to lack the capacity of recruiting the U2 snRNP to the branch point during the splicing reaction^78^. Both constructs were expressed for 5 hours in fully grown oocytes, where U2AF2 localized both in the nucleoplasm and in nuclear speckle-like compartments, whereas ΔRS-U2AF2 was only present in the diffuse phase of the nucleoplasm without any localization to nuclear speckles (Fig. 4P). These expression patterns were consistent with studies in human cells, where U2AF2 localizes to nuclear speckles and the removal of its RS domain disrupts this localization^79,80^. We then allowed oocytes to resume meiosis and monitored the timings of oocyte division in the two sets of oocytes. We found that the global percentage of oocytes undergoing division were comparable in both conditions (31/36 oocytes for WT U2AF2 versus 40/47 for ΔRS-U2AF2). However, only full-length U2AF2 expression had an impact on oocyte meiosis progression, causing a significant 4-hour delay in first polar body extrusion compared to the ΔRS-U2AF2-expressing oocytes (Fig. 4P-Q). This first oocyte division needs to be tightly regulated, and such significant delays in anaphase I progression along with first polar body extrusions are known to have major deleterious consequences on oocyte ploidy and capacity to turn into a viable embryo^81,82^. These data suggest that concentrations of spliceosome core proteins, such as U2AF2, should be fine-tuned in Prophase I to prevent meiotic defects, while showing that specific perturbations of even a single predicted splicing factor in fully grown oocytes can cause division phenotypes known to have deleterious consequences on oocyte fitness. Altogether, these experimental validations of selected resource-derived predictions reveal how both broad and specific splicing perturbations in fully grown oocytes can disrupt meiotic progression and, thus, the developmental potential of oocytes.

## Discussion

Decades of research established that pre-mRNA splicing occurs co-transcriptionally, while the nascent RNA remains covalently attached to RNA Polymerase II and, consequently, to the DNA template^83^. More recently, high-throughput sequencing has been an invaluable tool to visualize nascent transcripts, enabling observations indicating that co-transcriptional splicing efficiency was lower in mouse liver than in fly tissues and arguing for the presence of post-transcriptional splicing^84^. There is now accumulating evidence that post-transcriptional splicing constitutes a major regulatory mechanism to control the timing of gene expression^15^. In fully grown oocytes, RNA Polymerase II is depleted and leads to detection of only residual transcription^2–5^, which we show here to have no apparent role in correct oocyte division. By contrast, we demonstrate for the first time that splicing modulation at this stage has multiple deleterious effects on oocyte development. Specifically, the perturbations can impair maintenance of the nucleus in a central position, compromise the rapid assembly of the acentrosomal spindle, and disrupt timely cytokinesis and cell division. Our collective data thus indicate that a properly functional splicing machinery and transcription-independent splicing are necessary for oocytes to proceed through division in an accurate fashion and preserve their full developmental potential. Oocytes may use this post-transcriptional splicing as a mechanism to coordinate their cell cycle progression with the hormonal cues that regulate ovulation within ovarian follicles, thereby ensuring the timely expression of appropriate mRNA isoforms as they are finalizing preparations for meiosis resumption. In this developmental context, it is logical that three distinct splicing modulators converge on a shared set of genes implicated in the regulation of meiotic progression and division.

Recent investigations, mostly in human cell lines, revealed that RNA splicing can occur far from nuclear speckles, in their vicinity, or both, depending on transcript characteristics^19,21,22,85^. The localization of pre-mRNAs to nuclear speckles enhances splicing efficiency and lowers splicing entropy for a large subset of transcripts with specific features such as high GC content in exons and introns as well as shorter introns^19,21,85^. In fully grown oocytes, splicing activity occurs primarily in nuclear speckles, which are dissolved by Tubercidin^4^. In the case of this nuclear speckle dissolving compound for instance, our computational analyses indicate that an acute treatment impacts transcripts that contain a significantly higher GC content in both exons and their downstream introns (Fig. S4A, E, F), and significantly shorter intron lengths (Fig. S3A-B and S4B). These data are consistent with the splicing consequences of acute genetic dissolution of nuclear speckles recently shown in human colon-derived cell lines^85^, implying that nuclear speckles in fully grown mouse oocytes enable RNA processing via molecular means comparable to the ones described in human somatic cells. More broadly, these results underscore the utility of our resource to detect specific RNA splicing features potentially conserved across cells and species, with implications for broader research on biomolecular condensate and RNA biology.

Our study’s findings and resource have implications for fundamental and translational human fertility research. Indeed, multiple lines of evidence suggest the conservation of cellular and molecular events happening in both mouse and human oocytes at the end of their growth. For instance, fully grown mouse and human oocytes both present a centrally localized nucleus, a position that predicts successful oocyte development in both species^86,87^, and thus, their embryonic potential. In parallel, and similar to recent work suggesting the evolutionarily conservation of RNA Polymerase II degradation in mice and humans^5^, our preliminary evidence calls for a degree of conservation in nuclear speckle reorganization at the end of growth in both mouse and human oocytes. In a developmental context, nuclear speckle architecture in *Drosophila* and mouse oocytes depends on a mechano-transduction cascade driven by cytoskeletal forces^3,4,72^. In the context of aging, oocyte quality is also known to decrease with maternal age^88,89^, as well as both ovarian^90–92^ and oocyte stiffnesses^93^ are known to be affected by aging. The combined studies thus suggest that transcription-independent splicing activity may also contribute to the developmental potential of fully grown human oocytes, with clinical implications for in vitro fertilization efforts. Moreover, since cytoskeletal forces, and therefore mechano-transduction, are sensitive to external mechanical cues in both oocytes^94^ and somatic cells^95,96^, nuclear speckle organization and splicing may likewise be altered with maternal age in humans. Considering that both pharmacological and splice switching oligonucleotide-based corrections of splicing decisions are promising therapeutic strategies across pathologies^16,97–99^, this study provides a solid computational base for the future design of such strategies in fertility clinics.

## Acknowledgements

We thank all members of the Verlhac-Terret and Al Jord laboratories for discussions, the CIRB animal and Orion facilities. We also thank Sophie Bonnal and Juan Valcarcel for discussions and for providing the U2AF2 plasmids. The Verlhac-Terret laboratory is supported by CNRS, INSERM, Collège de France and the Bettencourt Schueller Foundation. MHV acknowledges the support of the Fondation pour la Recherche Médicale (DEQ201903007796 to MHV), of the Institut National du Cancer grant Inca-PREVBIO 2021-161 to MHV) and of the Fondation ARC pour la Recherche sur le Cancer (ARCPGA2023110007323_7953 to MHV). HAS was supported by Agence Nationale de la Recherche (ANR-18-CE13 to MHV). This work was supported by the France Génomique national infrastructure, funded as part of the “Investissements d’Avenir” program managed by the Agence Nationale de la Recherche (contract ANR-10-INBS-0009). The Al Jord Laboratory is supported by the Spanish Ministry of Science and Innovation through the Agencia Estatal de Investigación grant PID2024-157329NA-I00, the Centro de Excelencia Severo Ochoa (CEX2020-001049-S, MCIN/AEI /10.13039/501100011033 / FEDER, UE), the Generalitat de Catalunya CERCA programme, and the CRG Core Technologies programme.

## Author contributions

Conceptualization: MHV, AAJ, AGO, MI

Methodology: MHV, AAJ, AGO, MI, CB, EL

Investigation: HAS, CDS, AGO, SL, CB, EL

Visualization: HAS, AGO, JOT

Funding acquisition: MHV, AAJ

Project administration: MHV

Supervision: MHV, AAJ

Writing – original draft: MHV, AAJ, AGO, HAS

Writing – review & editing: MHV, AAJ, AGO, MET, MI

## Data, code, and materials availability

The data discussed in this publication have been deposited in NCBI’s Gene Expression Omnibus^100^ and are accessible through GEO Series accession number GSE320390. Computational analysis, custom scripts, statistical testing and data visualization were performed in R v4.5.2 and RStudio v2025.09.0. All the code, environments, libraries, software, the inclusion table (raw output from vast-tools) and tables containing all splicing events can be accessed via the GitHub repository (https://github.com/andresgordoortiz/oocyte_splicing_ssa_pladb_tub_2025). RNA-Seq samples and Supplementary Data 1 and 2 are stored on the Gene Expression Omnibus (GEO) with accession number GSE320390. We also provide the following interactive browser (*<u>Oocyte Splicing</u> <u>Explorer</u>*) to access and download desired data generated by this study.

## Methods

### Mouse oocyte collection

All animal studies were performed in accordance with the guidelines of the European Community and were approved by the French Ministry of Agriculture (authorization N°75– 20 1170) and by the Direction Générale de la Recherche et de l’Innovation (DGRI; GMO agreement number DUO-5291). Mice were housed in the animal facility on a 12-h light/dark cycle, with an ambient temperature of 22– 24°C and humidity of 40–50% and received food and water ad libitum. Mice used in this study include female C57BL/6J (Charles River Laboratories; 9 to 17 weeks old) and OF1 female mice (Charles River Laboratories; 7 to 17 weeks old). At least two mice were used per experiment. Ovaries were extracted from mice as previously described^101^ into M2 + Bovine Serum Albumin (BSA; Sigma, A3311) medium pre-warmed at 37°C and supplemented with 1 μM Milrinone (Merck, M4659) as in^102^ to maintain oocytes arrested in Prophase I. Ovarian follicles were punctured with surgical needles to release oocytes from antral 30 follicles (end of oocyte growth). All subsequent oocyte culture and live imaging steps were then carried out in M2+BSA+1 μM Milrinone under oil at 37°C (Sigma, M8410). For imaging without oil, µ-Slide 8 Well Glass Bottom ibidi plates were used.

### Immunofluorescence of human oocytes

Immature human oocytes, not arrested in meiosis II and thus not usable for patients, were collected, as approved by the local ethics committee of the Hospices Civils de Lyon (agreement number 22_5725). All patients gave informed consent. Patients undergoing Assisted Reproductive Technologies (ART) for intracytoplasmic sperm injection (ICSI) had multi-follicular ovarian stimulation. When the ovarian follicles were mature, patients underwent follicular fluid puncture to recover the cumulo-oocyte complexes (COCs). The follicular fluid was screened, and the COCs were cultured in media (GIVF, Vitrolife, Sweden) at 37°C, 5% CO2, and 5% O2 in an incubator during 1 h, and then enzymatically (80 IU/ml recombinant human hyaluronidase, ICSI Cumulase, Origio) and mechanically denuded. The denuded oocytes were selected under the light microscope. Only oocytes arrested in meiosis II were used for ICSI. The other oocytes in Prophase I, unsuitable for ICSI, were used in this study.

Human oocytes were fixed in 4 % paraformaldehyde at 37 °C for 30 min. They were permeabilized and pre-blocked for 2 hours in PBS with 0.1% Triton-X (Sigma, 93443) and 3% BSA (Sigma, A2153). Oocytes were extensively washed with PBS buffer between solutions. Oocytes were incubated with the primary antibody, mouse IgG1 anti-SRSF2/SC35 (1:200; ab11826, Abcam), during 2 hours at 4 °C and with the secondary anti-mouse antibody coupled to Alexa Fluor-488 (1:400; ThermoFisher) for 1 hour at room temperature. 30 minutes before reading under the spinning disk microscope (Andor Spinning disk, CIQLE platform), DAPI is added (1:600; Sigma, 10236276001). Primary and secondary antibodies were diluted in PBS with 0.1% Triton-X and 3% BSA. The focus was set on the germ vesicle, the z step was 1 µm, and 15 stacks were taken above and below.

### Live markers and drugs

Spy-DNA (Spirochrome, SPY650-DNA, TebuBio) and Spy-Tubulin (Spirochrome, SPY555-Tubulin, TebuBio) were diluted in anhydrous DMSO to make 1 mM stock solution kept at -20°C. They were used at 1 μM for 30 minutes prior imaging. The following inhibitors were used: DRB at 100 µM (D1916, Sigma-Aldrich), α-amanitin at 0.1 mg/mL (A2263, Sigma-Aldrich), Tubercidin at 10 μM (TO642, Sigma-Aldrich), Spliceostatin A at 0.1 and 1 µM (HY-16466, Clinisciences), Pladienolide B at 3 µM (T16551, Clinisciences). Dimethylsulfoxide (D2650, Sigma-Aldrich) was used as a solvent for all inhibitors before addition to M2 + BSA + 1 μM Milrinone medium. The culture medium containing live markers or drugs at their working concentration was freshly prepared and prewarmed at 37°C. Oocytes were washed three times in an inhibitor-supplemented medium and incubated for 5 hours before proceeding. For oocyte division experiments, oocytes were washed out of the inhibitors six times with M2 + BSA medium (without Milrinone) and allowed to proceed in the microscope chamber for up to 20 h.

### Microinjection

Oocytes were microinjected with cRNAs using an Eppendorf Femtojet microinjector. Oocytes were kept in Prophase I between 5 to 6 hours to allow expression of fusion proteins. Live imaging and oocyte isolation were carried out under oil at 37°C, while all incubations with drugs and ibid-plate imaging were carried out without oil at 37°C. We used the following constructs: H2B-RFP^42^, EB3-GFP^38^, Utr-GFP^42^, GFP-U2AF2 (this study), GFP-IBB-ΔRS-U2AF2 (this study). pRN3-GFP-U2AF2 and pRN3-GFP-IBB-ΔRS-U2AF2 were obtained by subcloning mouse U2AF2 and IBB-ΔRS-U2AF2 into BspEI/XhoI cloning site of a pRN3-GFP vector (GenScript). The Importin beta binding domain (IBB), which allows import of ΔRS-U2AF2 into the nucleus, comes from the Rango construct^37^. In vitro synthesis of capped cRNAs was performed as previously described^103^. cRNAs were centrifuged at 4°C for 45 min at 13,000 rpm before microinjection.

### Live imaging

Spinning-disk images were acquired at 37°C using a Plan-APO ×40/1.25 NA objective on a Leica DMI6000B microscope enclosed in a thermostatic chamber (Life Imaging Service) equipped with a Retiga 3 CCD camera (QImaging, Burnaby) coupled to a Sutter filter wheel (Roper Scientific) and a Yokogawa CSU-X1-M1 spinning-disk. Metamorph 40 software (Molecular Devices) was used to collect data.

### Image analysis and quantifications

All images were analyzed on Fiji (Version 2.16.0/1.54p).

Quantification of the time for nuclear envelope breakdown and polar body extrusion was calculated according to the difference between nuclear envelope breakdown timepoint and the beginning of the movie acquisition, and the difference between polar body extrusion and nuclear envelope breakdown timepoints, respectively (frames were converted to hours). Quantification of the durations of the different meiotic division phases (microtubule ball, spindle bipolarization, spindle migration, and polar body extrusion) was calculated according to the sequential timepoint differences between these phases (frames were converted to hours). Microtubules intensity was quantified using a defined ROI at different timepoints after background removal on Fiji. Quantification of nucleus position: ρ = d (nucleus center - oocyte center)/(oocyte radius - nucleus radius). ρ varies between 0 (perfectly centered nucleus) and 1 (completely off-centered nucleus). Cytoplasmic F-actin intensity was quantified averaging five different measurements of defined regions of interest (ROI) of 100×100 pixels on a single Z-slice and normalized by the nucleus intensity on that same Z-slice. Total nuclear SC35 signal intensities in human oocytes was quantified using z-projections covering the entire nucleus and normalized by the cytoplasmic signal intensity. To quantify number and size of nuclear speckles per oocyte, an SC35 signal threshold-based mask was created before extracting speckle number and area. Nuclear speckle volumes were inferred from the radii obtained through area measurements. Chromatin-associated speckles were identified based on chromatin-speckle contact, and the percentage of chromatin-associated speckles was calculated relative to the total number of nuclear speckles within each oocyte.

### Statistical analysis

All graphs and statistical analyses were generated using MS Excel (Version 16.78) and GraphPad Prism 10. SD was used as y-axis error bars for bar charts and curves plotted from the mean value of the data. Statistical significance based on Mann-Whitney test and Kolmogorov-Smirnov test was calculated in Graphpad Prism 10. All data are from at least two independent experiments. In the figures, significance is designated as follows: ****= p < 0.0001, ***= p < 0.001, **= p < 0.01, *= p < 0.05. Nonsignificant values are indicated as n.s.

### RNA sequencing

Fully grown oocytes from C57BL/6J strain were maintained in Prophase I with μM Milrinone and treated for 5h either with DMSO (control) or with 10 μM Tubercidin, 1 μM Spliceostatin A or 3 μM Pladienolide B. They were then washed in PBS. Groups of 15 oocytes were collected in triplicates. Individual samples were collected in 2 µl PBS then resuspended in lysis buffer supplemented with 1% of β-Mercaptoethanol, frozen in liquid nitrogen and conserved at -80°C overnight. Total RNA was extracted using the RNAqueous Micro Total RNA Isolation Kit (Thermofisher, AM1931). After unfreezing, RNA extraction was carried out following the manufacturer’s instructions. RNA was eluted twice in 10 μl elution buffer. Samples were then treated with DNAse I and stored at -80°C. Oocytes cDNA libraries and RNA sequencing Library preparation and Illumina sequencing were performed at the Ecole normale supérieure GenomiqueENS core facility (Paris, France). 10 ng of total RNA were amplified and converted to cDNA using SMART-Seq mRNA kit (Takara). Afterwards, an average of 500 pg of amplified cDNA was used to prepare the library following Nextera XT DNA kit (Illumina). Libraries were multiplexed by 15 on P4 flow cells. A 68 bp read sequencing was performed on a NextSeq 2000 device (Illumina).

The analyses were performed using the Eoulsan pipeline^104^, including read filtering, mapping, alignment filtering, read quantification, normalisation and differential analysis: Before mapping, poly N read tails were trimmed, reads ≤40 bases were removed, and reads with quality mean ≤30 were discarded, yielding an average of 360 million reads per biological replicate. The analysis included 15 samples across 4 experimental conditions: control (n=6), tubercidin (n=3), pladienolide B (n=3), and spliceostatin A (n=3). Since pladienolide B samples were sequenced in a different day, 3 extra control replicates were accounted to avoid batch effects and used exclusively for the Pladienolide B analysis. Reads were then aligned against the *Mus musculus* genome GRCm38 assembly (GCF_000001635.20 NCBI RefSeq) using the default configuration of the nf-core/rnaseq v3.22.2^105^, and gene level quantification was assessed with Salmon as the pseudoaligner.

### Gene Expression & RNA Splicing Analyses

The sample counts were normalized using DESeq2 1.8.1^106^. P-values were adjusted for multiple testing using the Benjamini-Hochberg false discovery rate (FDR) method. Genes were considered differentially expressed if they met an adjusted p-value <= 0.05 and an absolute log2Fold Change (log2FC, Treatment/Control) >= 1.5. We noticed that some genes produced unrealistic log2FCs—over 20 in some cases—due to high sample dispersion and consistent low read counts in control samples. To address this issue, we applied log2 fold change shrinkage using the apeglm method^107^, which has been shown to provide more accurate effect size estimates than other shrinkage methods. Both shrunken and unshrunken log2 fold changes were retained for comparison and both yielded a similar number of differentially expressed genes (see Supplementary).

For mRNA splicing analyses, event level identification was performed using vast-tools v2.5.1^52^. Briefly, trimmed samples were aligned to the mm10 assembly of VastDB using vast align and outputs were merged using vast combine. A custom Nextflow pipeline to replicate our analysis can be accessed on this link (https://github.com/andresgordoortiz/vast-tools_nextflow). The output inclusion table contains all identifiable splicing events across samples. Differential splicing quantification was then assessed using betAS v1.2.1^53^, a tool that models inclusion proportions using beta distributions based on RNA sequencing reads supporting inclusion and exclusion of splicing sequences. First, events with less than 10 supporting reads in at least one sample were filtered. To confirm that this threshold yielded robust results, we repeated event detection across stricter minimum read-support thresholds, up to 100 reads. The total number of quantifiable events decreased smoothly and predictably with increasing stringency (from 35,375–78,777 events at 10 reads to 6,411–31,164 events at 100 reads, across event types and conditions), with no abrupt drops that would suggest an unstable threshold. This filtering choice does not, however, reduce confidence in individual significant calls, since betAS estimates each event’s false-positive rate from beta distributions fitted to its own read counts; read-depth uncertainty is therefore already built into the significance test itself, rather than relying on an arbitrary fixed coverage cutoff. Then, beta distributions were fitted for each splicing event, with inclusion rates and quality scores serving as surrogates for the two shape parameters, yielding distributions centred on the percent spliced-in (PSI) values with precision reflecting read coverage. For each event, the null distribution of PSI differences was estimated through random sampling from beta distributions fitted to each sample and the false-positive rate (FPR) was calculated as the proportion of 1,000 random simulations yielding ΔPSI values equal to or more extreme than the empirically observed ΔPSI between groups. Events were considered significantly differentially spliced if they met two criteria: FPR <= 0.05 and an absolute ΔPSI >= 10%, where ΔPSI represents the difference in mean PSI values between the treated and control samples. To test for any directional bias of splicing changes, we classified every covered event by the sign of its mean PSI difference across the same range of read-support thresholds (10–100 reads). This revealed small skews for some drugs (Pladienolide B and Spliceostatin A toward retention, ∼51–52%; Tubercidin toward excision, ∼48.7–49.1%) that reached statistical significance given the large sample sizes but deviated from a 50/50 split by only 1 to 2 percentage points, indicating that the overall distribution of splicing changes was indeed symmetrical. The predicted functional impact of significant splicing events was obtained from the VastDB database v3 of the mm10 build (https://vastdb.crg.eu/downloads/mm10/PROT_IMPACT-mm10-v3.tab.gz).

### Sequence Features Analysis

Analysis of splice site strengths, exon- and intron-related features, and other features related to the spliced and neighboring sequences (e.g. length, GC content, splice site predicted strength, branch point analysis) was performed using the Matt toolkit (Gohr, André, and Manuel Irimia. ‘Matt: Unix Tools for Alternative Splicing Analysis’) through matt cmpr_exons or matt cmpr_introns with the -notrbts flag. Only significant exons or introns were used as the input, alongside a random sample of 10,000 non-significant events (under any treatment) as the background control. A Docker container with our Matt installation can be found on this link (https://hub.docker.com/repository/docker/andresgordoortiz/matt-container/tags/latest/sha256:6b729b82e312f06153f8faeb643402df4e1729485fba90359a6cf3928c24c2 0e). For every feature analyzed, groups in every treatment were statistically compared to the background control using Mann-Whitney tests.

### RBPs Motif Enrichment Analysis

The rMAPS2 server (http://rmaps.cecsresearch.org/)^64^ was used to explore the RNA-binding protein (RBPs) motif enrichment in the differentially spliced exons, introns, alternative 5’ splice sites (A5SS), and alternative 3’ splice sites (A3SS). For each splicing event type, rMAPS2 scanned for occurrences of over 110 known RBP consensus motifs within these genetic positions: upstream exon, upstream flanking intron, target exon/intron, downstream flanking intron, and downstream exon regions. Motif density scores were calculated using a sliding window of 50bp for exons and 250bp for introns and represented as the overall percentage of nucleotides covered by each motif. rMAPS2 calculates statistical significance of motif enrichment using Mann-Whitney tests comparing upregulated versus background and downregulated versus background events at each sliding window within each genetic position. Background events are the non-significant splicing events in each treatment. Since statistical significance varies alongside the sliding window, rMAPS2 output consists on a table containing the lowest (most significant) p-value for a given motif for every genetic position. For simplicity and interpretability, only the most significant motif for each RBP and genetic position was selected for display.

### Gene Ontology Analysis

Gene Ontology (GO) analysis of the genes containing significantly spliced events was performed using enrichGO from clusterProfiler^108^, using the org.Mm.eg.db package as the organism database and universe, and setting the adjusted p-value [Benjamini-Hochberg] cut-off to 0.05. GO terms from the 3 categories—Biological Process, Cellular Component and Molecular Function—were sorted by their number of genes.

## Supplementary Figure Legends

**Supplementary Figure 1. Nuclear speckle evolution at the end of human oocyte growth**

Human oocytes arrested in Prophase I after follicle aspiration and were therefore unsuitable for in vitro fertilization. The oocytes were classified according to their progressive degree of chromatin condensation^4,109^ (NSN, Trans, and SN).

**A.** Immunofluorescence images of fixed NSN and SN human oocytes labelled for nuclear speckles (anti-SC35) and chromosomes (DAPI). Maximal projections of 4 Z-steps are shown. Scale bar 10 µm.

**B.** The number of nuclear speckles decreases at the end of human oocyte growth. N = 8 NSN human oocytes; N =7 Trans human oocytes; N = 15 SN human oocytes. Violin plots with median ± quartiles. P values derived from a Kruskal-Wallis test; ** = p<0.005.

**C.** The volume of nuclear speckles increases at the end of human oocyte growth. Violin plots with median ± quartiles. P values derived from a Kruskal-Wallis test; * = p<0.03.

**D.** Total SC35 signal intensity per nucleus does not change across stages, similar to what was described in mouse oocytes. Violin plots with median ± quartiles. P values derived from a Kruskal-Wallis test; ns = p>0.05.

**E.** The sum of nuclear speckle volumes per nucleus decreases only in SN oocytes, suggesting dissolution of a subpopulation of speckles in SN oocytes which would be consistent with RNA Polymerase II degradation and a transcriptional halt at this stage. Violin plots with median ± quartiles. P values derived from Mann-Whitney tests; ns = p>0.7; * = p<0.05.

**Supplementary Figure 2. Transcriptional inhibition by α-amanitin of fully grown oocytes does not impact subsequent meiosis progression.**

Percentage of control and α-amanitin-treated oocytes that undergo NEBD and extrude their first polar body (PBE) as a function of time (h) after washout from drugs. The 0h time point marks the first time point after washout. N= 43 oocytes for control, N= 58 oocytes for α-amanitin, 3 independent experiments.

**Supplementary Figure 3. Splicing modulators disrupt mRNA splicing in fully grown oocytes**

**A.** Volcano plot showing splicing events by their spliced-in rate (PSI) and the False Positive Rate (FPR) for each treatment. Significant splicing events reach a |ΔPSI| of at least 0.1 (10%) and a FPR lower than 0.05, in which case they are colored by their corresponding treatment. The number of significant events is displayed.

**B-C.** Heatmaps showing splicing changes of significant exons (B) and introns (C) across different treatments. Tiles represent row-wise z-scores of PSI values for significant splicing events (FPR ≤ 0.05 and |ΔPSI| ≥ 0.10) when comparing each drug treatment to the control. Spearman correlations (within-group and between-group) are shown below each heatmap. Low positive correlation coefficients within groups indicate low interdependence among replicates, whereas higher negative correlations between groups suggest opposite splicing behaviors (i.e., exons that are included in controls tend to be skipped in treatments, and vice versa).

**Supplementary Figure 4. Significantly spliced exons showcase sequence features related to splicing efficiency.**

**A-F.** Panels showing the transcript sequence features of significantly spliced exons across treatments. Exons are categorized as “Included” (increased inclusion, ΔPSI > 0.1), “Skipped” (increased skipping, ΔPSI < -0.1), or “Unchanged” (non-significant; randomly sampled n = 10,000 per treatment). Box plots display median and interquartile ranges. Statistical pairwise comparisons were performed using Mann-Whitney tests (*P ≤ 0.05, **P ≤ 0.01, ***P ≤ 0.001, ****P ≤ 0.0001). Outliers exceeding the 75th percentile were clipped for visualization purposes only; statistical tests and descriptors are shown in complete, unclipped data.

**Supplementary Figure 5. Significantly spliced introns showcase sequence features related to splicing efficiency.**

**A-F.** Panels showing the transcript sequence features of significantly spliced introns across treatments. Introns are categorized as “Retained” (increased retention, ΔPSI >= 0.1), “Excised” (increased excision, DPSI =< -0.1), or “Unchanged” (non-significant; randomly sampled n = 10,000 per treatment). Box plots display median and interquartile ranges. Statistical pairwise comparisons were performed using Mann-Whitney tests (*P ≤ 0.05, **P ≤ 0.01, ***P ≤ 0.001, ****P ≤ 0.0001). For length analysis, outliers exceeding the 95th percentile were clipped for visualization purposes only; statistical tests were performed on complete, unclipped data.

**Supplementary Figure 6. Significantly spliced exons and introns showcase sequence motifs functionally linked to splicing RBPs.**

**A.** Tracks showing the motif enrichment significance of RBPs on the neighboring sequences of included and skipped exons. Motif enrichment was calculated using rMAPS2, using the significant exon sequences as input, and all non-significant sequences as the background control. The positions correspond, from left to right (5’—>3’), to distributions—not precise locations—encompassing the last 50bp of the upstream exon, the first and last 250bp of the upstream intron, the first and last 50bp of the spliced exon and likewise for the downstream intron and downstream exon. Only the top 15 RBPs— according to lowest p-value—per splicing event and position were selected, and only the top 3 is labelled.

**B.** Comparison of RBP enrichment across genomic regions and drug treatments. Box plots display the distribution of -log10(p-values) for RNA-binding protein enrichment at five genomic positions across treatments and splicing outcome. Statistical significance between included and skipped exons within each region and treatment was assessed using the Mann-Whitney test (*P ≤ 0.05, **P ≤ 0.01, ***P ≤ 0.001, ****P ≤ 0.0001). Box boundaries represent the interquartile range (IQR), with the center line indicating the median. The positions correspond, from left to right (5’—>3’), to the last 50bp of the upstream exon, the first and last 250bp of the upstream intron, the first and last 50bp of the spliced exon and likewise for the downstream intron and downstream exon.

**C.** Tracks showing the motif enrichment significance of RBPs on the neighboring sequences of included and skipped introns. Motif enrichment was calculated using rMAPS2, using the significant intron sequences as input, and all non-significant sequences as the background control. The positions correspond, from left to right (5’—>3’), to distributions—not precise locations—encompassing the first and last 50bp of the upstream exon, the spliced intron and the first and last 50bp of the downstream exon. Only the top 15 RBPs—according to lowest p-value—per splicing event and position were selected, and only the top 3 is labelled.

**D.** Comprehensive RBP motif enrichment analysis across three splicing-modulating drugs of the significant introns or adjacent sequences. The heatmap shows the highest -log10 (p-values) for the top RBPs, sorted by maximum significance across conditions. Rows are clustered by Euclidean distance; columns are ordered by genomic position, drug and splicing direction (Retention, Excision). The positions correspond, from left to right (5’—>3’), to the first and last 50bp of the upstream exon, the spliced intron and the first and last 50bp of the downstream exon.

**E.** Comparison of RBP enrichment across genomic regions and drug treatments. Box plots display the distribution of -log10(p-values) for RNA-binding protein enrichment at five genomic positions across treatments and splicing outcome. Statistical significance between retained and excised introns within each region and treatment was assessed using the Mann-Whitney test (*P ≤ 0.05, **P ≤ 0.01, ***P ≤ 0.001, ****P ≤ 0.0001). Box boundaries represent the interquartile range (IQR), with the center line indicating the median. The positions correspond, from left to right (5’—>3’), to the first and last 50bp of the upstream exon, the spliced intron and the first and last 50bp of the downstream exon.

**Supplementary Table 1.** Gene expression table following treatment with the three splicing modulators.

**Supplementary Table 2.** RNA splicing analysis table following treatment with the three splicing modulators.

**Supplementary Table 3.** Prediction table of the impact of RNA splicing events caused by the three splicing modulators.

**Supplementary Table 4.** Gene ontology analyses (biological process, cellular component, molecular function) of genes containing significantly spliced events caused by the three splicing modulators.

**Supplementary Data 1.** Sequence-specific structural features of the exons, introns, and flanking sequences significantly affected by Tubercidin, Pladienolide B, and Spliceostatin A treatment, related to Figure 2G and Figures S4–S5. Features were extracted using the Matt toolkit and include maximum-entropy-based 5′/3′ splice-site strength scores, predicted branch point positions, polypyrimidine tract length, GC content, and neighboring intron/exon length, computed for each significantly spliced event (included/skipped exons and retained/excised introns) across the three treatments. Stored on GEO # GSE320390.

**Supplementary Data 2.** RNA-binding protein motif enrichment analysis of the exons and introns significantly affected by Tubercidin, Pladienolide B, and Spliceostatin A treatment, related to Figure 2H and Figure S6. Enrichment was computed with rMAPS2 across exonic and adjacent intronic sequences, reporting enrichment scores and significance for RBP binding motifs and splicing regulators separately for included/skipped exons and retained/excised introns, for each treatment. SF3B1 motif was manually included in the dataset from ENCORI. Stored on GEO # GSE320390.

## Legends to Movies

**Movie S1: Control oocyte undergoing division after 5h in 1 μM Milrinone.** The oocyte is labeled for chromosomes (pink) and microtubules (green). 0h marks NEBD. Time is indicated in hours on the left corner. Movie related to Fig. 1C (top panel).

**Movie S2: DRB-treated oocyte undergoing division after 5h in 1 μM Milrinone + DRB.** The oocyte is labeled for chromosomes (pink) and microtubules (green). 0h marks NEBD. Time is indicated in hours on the left corner. Movie related to Fig. 1C (bottom panel).

**Movie S3: Control oocyte undergoing division after 5h in 1 μM Milrinone.** The oocyte is expressing H2B-RFP for chromosomes (pink) and EB3-GFP for microtubules (green). 0h marks NEBD. Time is indicated in hours on the left corner. Movie related to Fig. 4E (top panel).

**Movie S4: Tubercidin-treated oocyte undergoing division after 5h in 1 μM Milrinone + Tubercidin.** The oocyte is expressing H2B-RFP for chromosomes (pink) and EB3-GFP for microtubules (green).0h marks NEBD. Time is indicated in hours on the left corner. Movie related to Fig. 4E (bottom panel).

**Movie S5: Control oocyte undergoing division after 5h in 1 μM Milrinone.** The oocyte is expressing EB3-GFP for microtubules (green). 0h marks NEBD. Time is indicated in hours on the left corner. Movie related to Fig. 4E.

**Movie S6: Tubercidin-treated oocyte undergoing division after 5h in 1 μM Milrinone + Tubercidin.** The oocyte is expressing EB3-GFP for microtubules (green). 0h marks NEBD. Time is indicated in hours on the left corner. Movie related to Fig. 4E.

**Movie S7: Control oocyte followed during the 5h incubation in 1 μM Milrinone.** Movie related to Fig. 4L (top panel).

**Movie S8: Spliceostatin A-treated oocyte followed during the 5h incubation in 1 μM Milrinone + 1 μM SSA.** Time is indicated in hours on the left corner. Movie related to Fig. 4L (bottom panel).

