## Supplementary figures and images for "The post-transcriptional RNA splicing landscape driving oocyte division"

### Figure S1

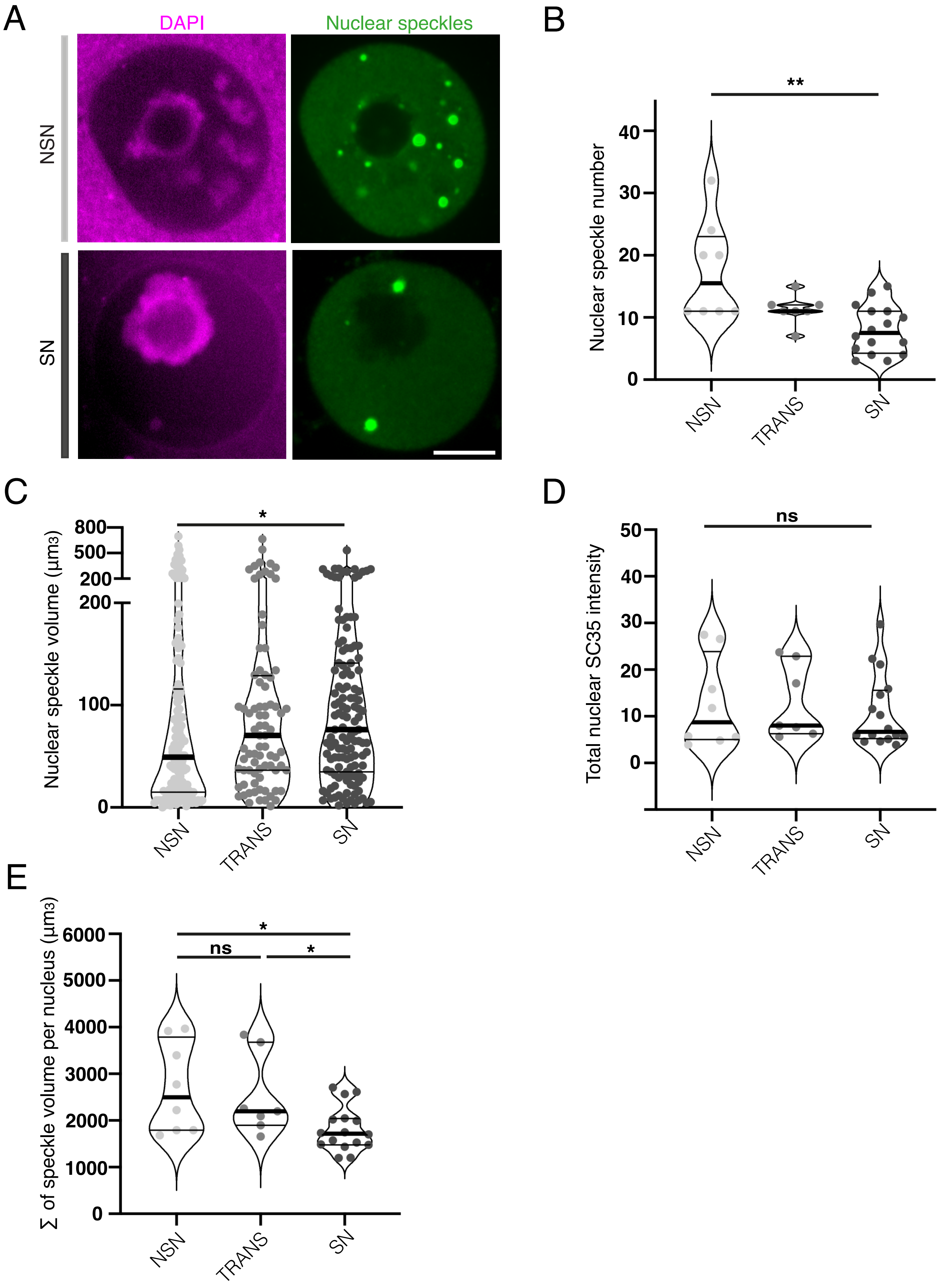

### Figure S2

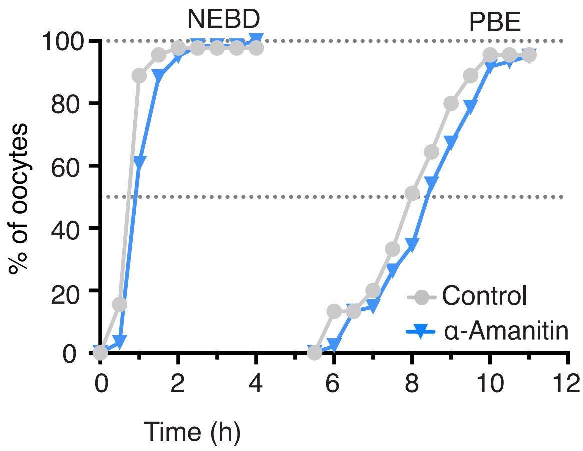

### Figure S3

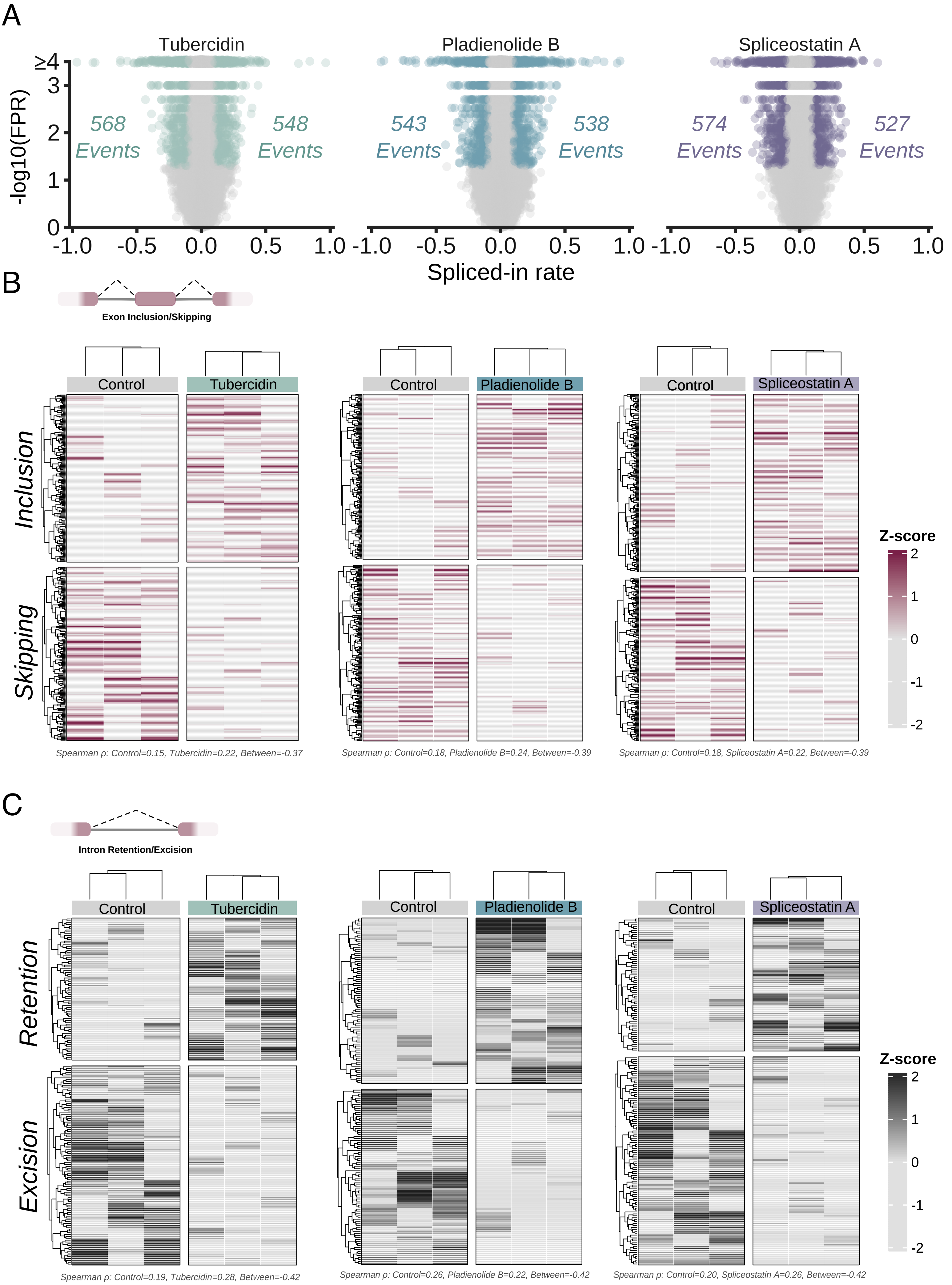

### Figure S4

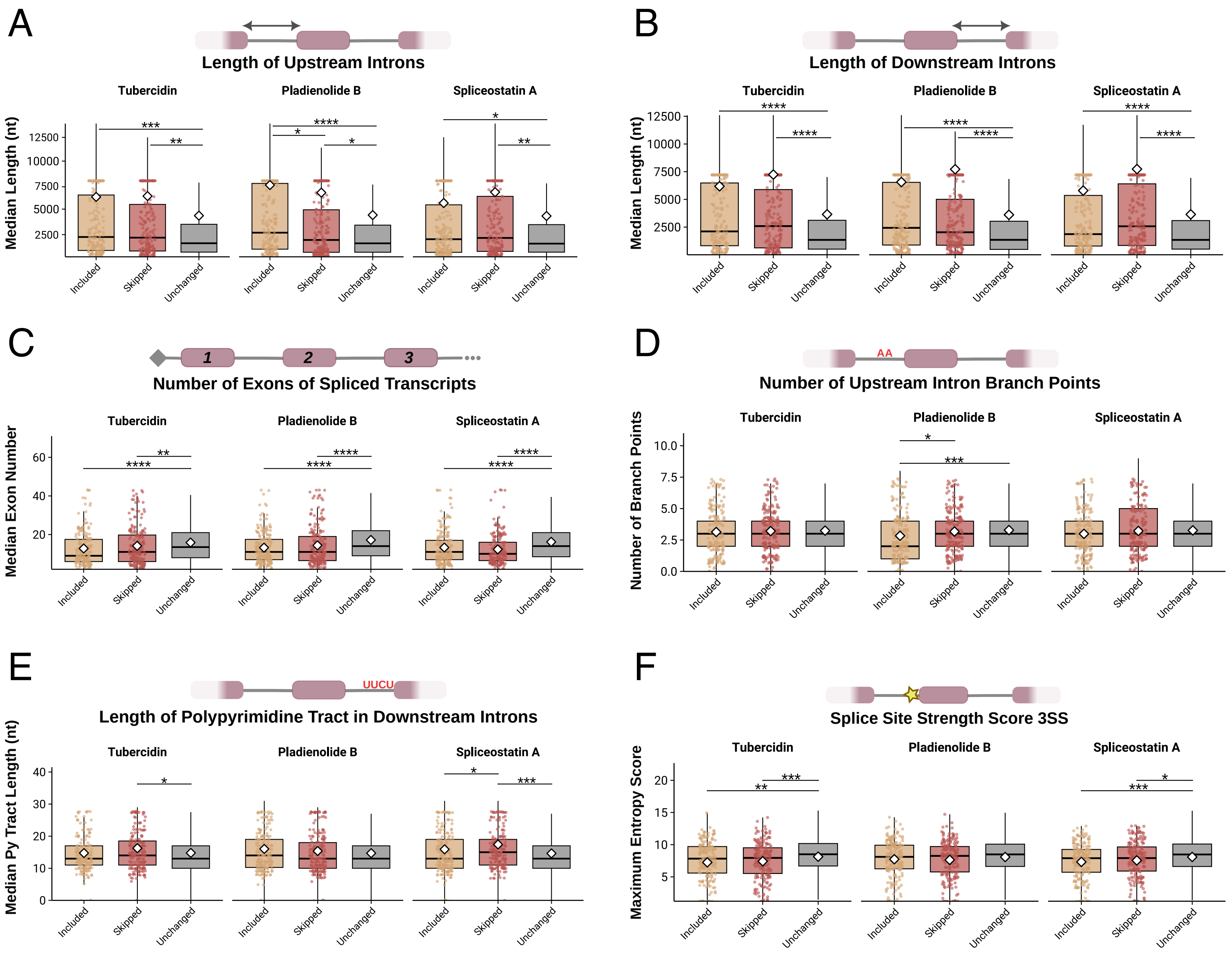

### Figure S5

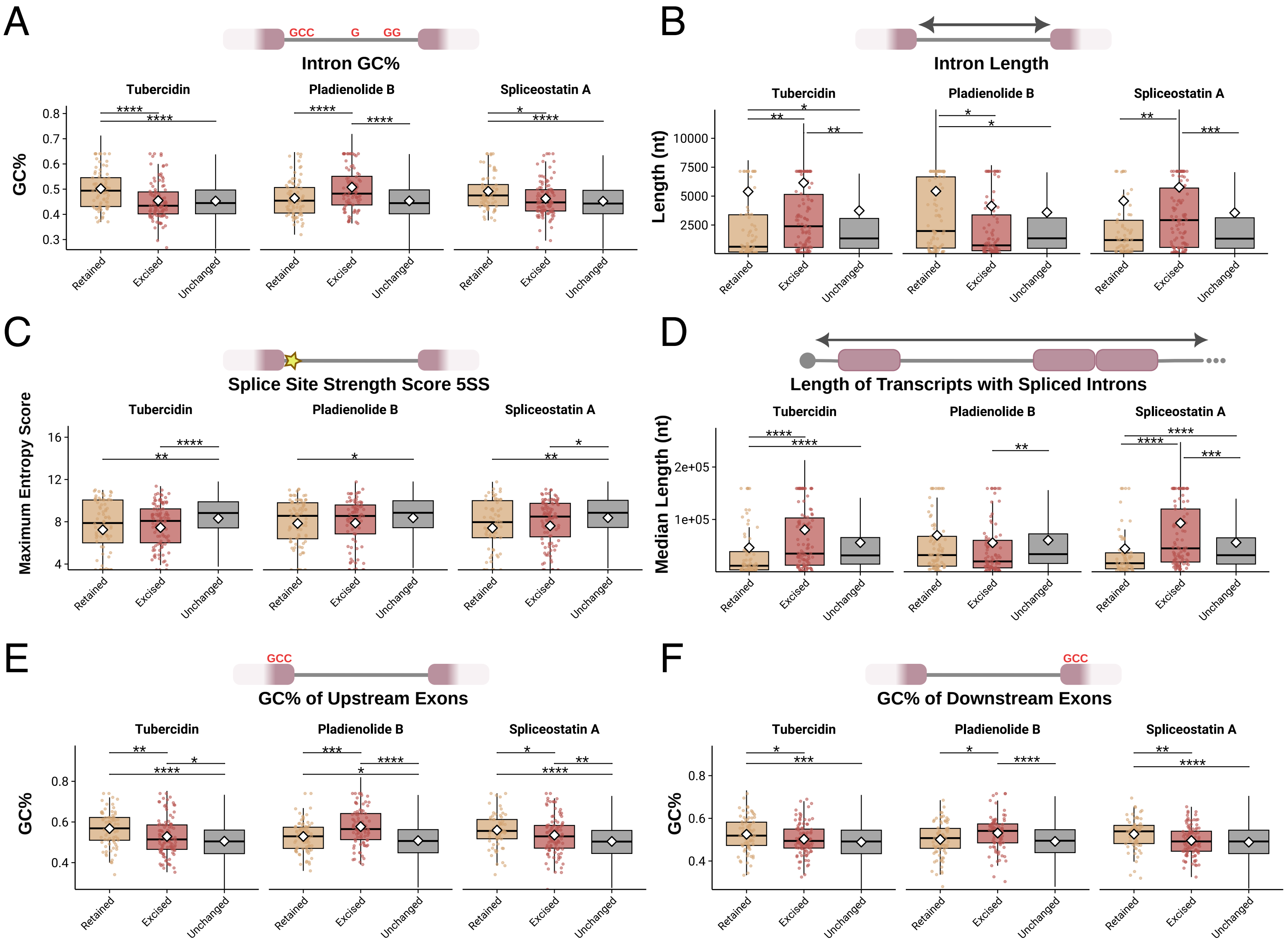

### Figure S6

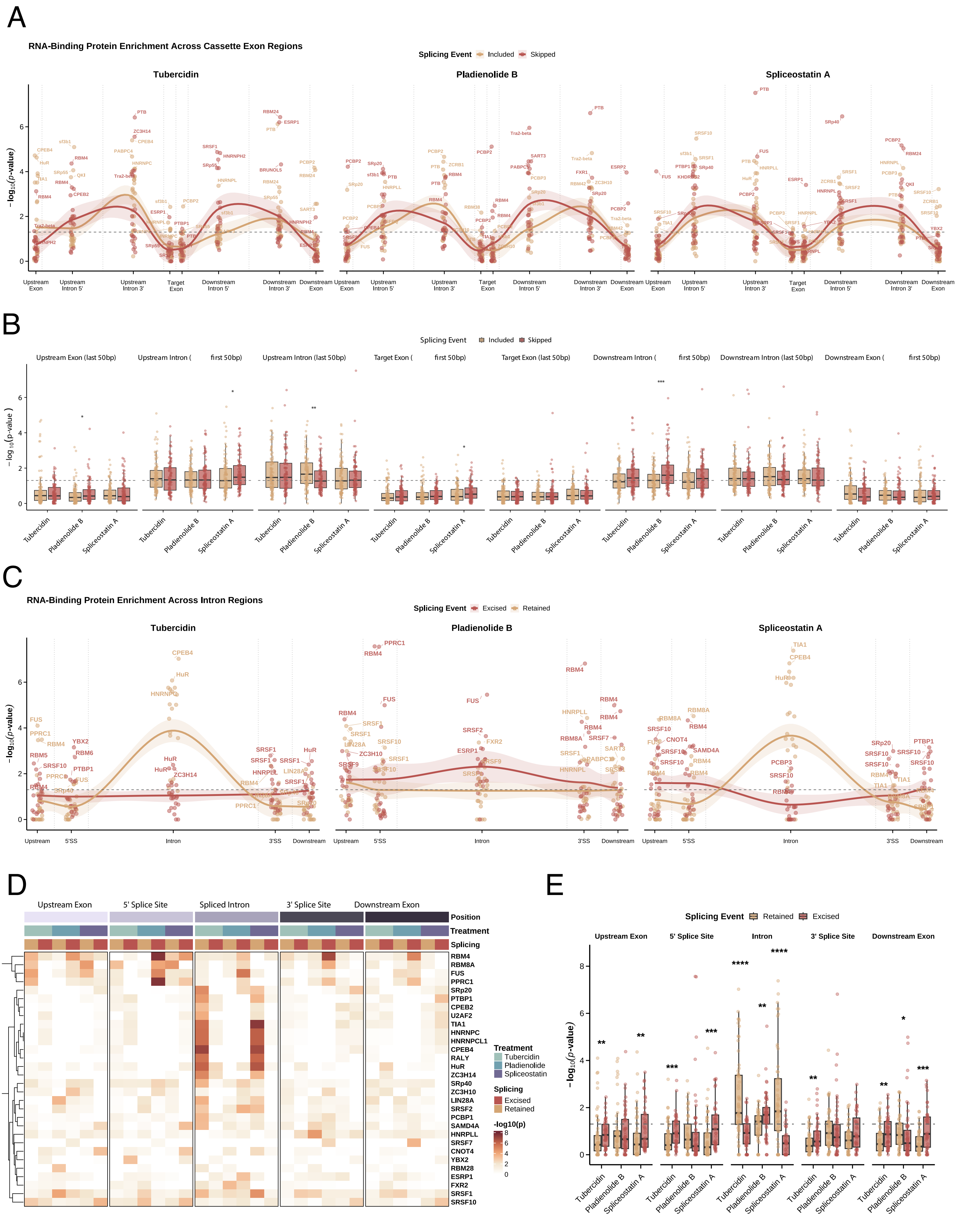
